# OCcAM: A tool for rapidly assessing impacts of offshore wind farms on seabirds

**DOI:** 10.1101/2025.08.22.671688

**Authors:** Gillian Clare Vallejo, James Robbins, Emily Nelson, Graeme Cook, Jonathan Abbatt

## Abstract

Offshore wind energy generation has a key role in reducing reliance on fossil fuels and mitigating climate change. However, it also has the potential to negatively impact the environment, including seabird populations. Quantifying and minimising impacts of wind projects on seabirds is necessary for ensuring compliance with environmental legislation and safeguarding populations. However, the timescales and complexity involved in assessment processes can limit the extent to which estimates can be leveraged towards balancing environmental impact and energy generation.

OCcAM is a simple, user-friendly and transparent industry-developed tool allowing rapid assessment of impacts on seabirds from offshore wind developments. Mortality rates of five seabird species are estimated using two UK industry-standard models: the Band collision risk model which predicts fatal collisions with turbine blades, and the matrix-based displacement approach that predicts mortalities associated with distributional responses to the presence of a wind farm. The tool supports simultaneous analysis of up to three input parameter sets and predictions can be expressed as a percentage of a focal population.

OCcAM allows auditable analyses to be run quickly and easily with no requirement for specific technical expertise, making it accessible to all stakeholders. It presents a variety of opportunities to facilitate ornithological assessment at strategic to project-specific scales. Here we demonstrate three potential applications of OCcAM: (1) predicting the cumulative impact of Scottish offshore wind farms upon protected seabird populations, (2) updating fatality estimates calculated for worst-case consented values using as-built input parameters and (3) optioneering of project design parameters relative to seabird risk.

## 1 Introduction

With the ever-increasing threat of climate change, expediting the transition to renewable energy production has never been more important. This is reflected in ambitious targets for global renewable energy production (Bolton, 2025; EU, 2023; COP28 *et al*., 2023). In order to achieve these targets, it is beneficial for governments to set out high level plans including an assessment of how the proposed growth in renewable energy will impact upon protected wildlife features, in line with environmental legislation (Dalal-Clayton and Scott-Brown, 2024).

Offshore wind energy has great potential for economically viable renewable energy generation in many countries and is expected to make a large contribution to meeting overall renewable energy goals (Ramirez *et al*., 2020). However, along with environmental benefits associated with offshore wind energy generation, there is also the potential to impact negatively upon some environmental receptors (Galparsoro *et al*., 2022).

Seabirds represent a key environmental receptor for offshore wind with three main mechanisms by which their populations may be directly affected by development (Drewitt and Langston, 2006). These are: 1) collision with the rotating blades of the wind turbines causing direct mortality; 2) displacement from habitat that would otherwise have been utilised, for example, from feeding grounds to habitat elsewhere that may be lower quality or more difficult to reach; and, 3) barrier effects, in which birds travel further when commuting between habitats, as a result of avoiding wind farms (Drewitt and Langston, 2006). The latter two, though not directly causing mortality, will result in increased energetic costs which may lead to reductions in survival and/or reproductive output.

The Environmental Impact Assessment (EIA) process should expediate development of offshore wind whilst protecting seabird populations, through identification, avoidance, reduction and offsetting of predicted population level risks (Morgan, 2012). This can be achieved if the EIA process is supported by tools that allow rapid and streamlined indicative assessments which are transparent and readily accessible to a range of end users for informing decision-making. This might relate to predicting potential impacts of future developments in different areas based on existing data or allowing reassessment of estimated impacts of existing developments, where build-out parameters may differ from consented maximum design scenarios. Such tools could equally inform advice provided by regulators and their advisors during project development, and support decisions made by developers to modify their design parameters to avoid adverse effects on seabirds. Such actions are difficult to achieve with the tools currently available to the industry, since the assessment pathway involves combining outputs from multiple discrete steps, some of which can be complex, time-consuming, computationally intensive and/or require specialist knowledge to apply.

OCcAM (Ornithology Cumulative Assessment Model) is an Excel-based seabird impact assessment tool which has been developed to fill this gap. The tool is structured as a simple spreadsheet which can be used to derive estimates of mortality associated with collision and distributional responses (displacement and barrier effects). Additionally, the tool allows these impacts to be apportioned to a focal population of interest and can apply corrections to account for the mismatch in “currency” between estimates of impacts on site, which generally relate to all birds, and population size estimates, which tend to be focused on breeding adults. Mortalities can thereby be attributed to, for example, a regional population, as required under the EIA process, as well as to specific features of protected sites such as breeding adult populations at Special Protection Areas (SPAs), required for UK and EU Habitat Regulations Assessment (HRA).

Here we describe the development, structure and usage of the OCcAM tool illustrated using three case studies representing alternative use-cases.

## 2 Tool conception

At a strategic level, a key activity for successful growth of offshore wind energy capacity in line with environmental legislation is strategic planning by asset managers and regulatory bodies, involving broad-scale EIA to ensure that environmental receptors are not adversely affected. As part of this, it is necessary to determine the cumulative impacts of current and planned offshore wind energy developments on seabird populations and, where there is potential to adversely affect seabird populations protected under the Habitats Regulations, develop strategic level plans to implement compensatory measures that can offset these impacts. This process is currently underway in the UK (DEFRA, 2025a; Tapia-Harris and Evans, 2024).

However, it is difficult to calculate meaningful estimates of cumulative effects, even for existing projects, due to project-specific differences in methods and parameters used for the project-only assessments upon which cumulative assessments are often based, together with changes in statutory advice for undertaking assessments over time. Methodologies currently being employed to attempt to address this problem can lack transparency on underlying assumptions used and can take a long time to implement due to the complexities in pulling together all the different data required and standardising it as best possible (O’Brien, 2025).

Due to these complexities, a need was identified for a simple and quick approach to derive indicative cumulative impact estimates based on more readily available input parameters, to provide meaningful answers at a large scale. For high-level assessments, it should be sufficient to derive outputs that can give order of magnitude indications of likely effects on seabird populations, facilitating rapid decision making. It was considered that the methodologies applied should be straightforward to audit and update as new data and knowledge emerge.

OCcAM was therefore developed as an independent method to indicate the likely magnitude of cumulative seabird mortalities from present and future offshore wind energy developments in a cost-effective, rapid and transparent way. The methodology applied within OCcAM is based upon industry-standard assessment tools that have been applied in the UK and elsewhere in Europe for assessment of impacts of offshore wind on seabirds during the EIA process. OCcAM currently focuses on five key seabird species but could be adapted to incorporate other species as required. Species currently included are: black-legged kittiwake (*Rissa tridactyla*), common guillemot (*Uria aalge*), razorbill (*Alca torda*), Atlantic puffin (*Fratercula arctica*) and northern gannet (*Morus bassanus*). These were chosen as they are often identified as key species of concern in UK assessments.

Whilst the aim of the tool was to generate cumulative impact predictions, OCcAM also provides a solution to many other key challenges relating to the development of offshore wind energy. Two examples of such use-cases are presented in Sections 6 and 7 below.

## 3 Tool methods

### 3.1 Collision risk modelling

OCcAM predicts annual collision mortality for kittiwake and gannet only, since guillemot, razorbill and puffin generally fly below rotor-swept height for offshore wind turbines and are therefore generally considered to be at low risk of collision (Bradbury *et al*., 2014). Predictions are made using the Band (2012) collision model. This model predicts collision rates for a proposed offshore wind farm as a function of number of turbines, wind turbine specifications, expected time that the turbines will be operational, and the density of seabirds at the site, their flight heights and species-specific biometric and behavioural data. Predicted collision mortality in the absence of any bird avoidance behaviour is calculated as the product of three components: (1) the predicted number of rotor transits per month per species, based on bird densities, flight speeds and flight heights relative to rotor height, (2) the individual probability of a bird colliding with a turbine blade given that it flies through rotor-swept space, based on characteristics of the bird species and of the turbine rotors and (3) the proportion of time that turbines are expected to be operational (Band, 2012). The resulting value is corrected using species-specific avoidance rates that represent the avoidance action expected to be taken by the birds, giving a final predicted monthly number of individuals per species expected to collide with the turbine rotors.

The Band (2012) model has been used extensively in the UK, and can be implemented using a publicly available macro-enabled Excel tool (BTO, 2013). The OCcAM tool draws heavily on this work and this spreadsheet forms the basis for the collision risk calculations applied within OCcAM.

The Band (2012) model also forms the basis of the stochastic collision risk model, known as sCRM (Caneco *et al*., 2022), which is the current recommended method for use in UK EIA through guidance from statutory advisors (SNCBs, 2024). The sCRM builds on the original model by allowing incorporation of variability and uncertainty in the input parameters, randomly sampling from distributions of input parameters over iterative model runs to derive output distributions. Since OCcAM was developed as a simple and transparent tool to provide instantaneous predictions it does not provide estimates of uncertainty. However, it’s predictions will be very close to the average values generated by the stochastic model if run with the same input parameters.

More detailed information regarding the Band model, including assumptions and caveats can be found in Band (2012).

### 3.2 Distributional responses modelling

Mortalities arising from distributional responses can be predicted for all five species within OCcAM though there is an option to suppress this for kittiwake since kittiwake is sometimes considered to be at low risk from displacement (SNCBs, 2022). The method used by OCcAM to predict these mortalities is known as the “matrix” approach (SNCBs, 2022). This method is currently the predominant recommended approach for use in UK EIA through guidance from statutory advisors (NatureScot, 2023a; Parker *et al*., 2025). The method uses mean seasonal-peak abundances of birds predicted to be either using or passing through the array area and surrounding species-specific displacement buffer (currently recommended to be 2km for most species). This is assumed to represent the population of birds with the potential to experience displacement effects. Predicted mortalities are calculated as the product of the abundance and species-specific rates representing (a) susceptibility to displacement and (b) mortality rates of displaced individuals. The matrix approach is so-called because results are generally provided within a matrix representing a range of possible displacement rates along one axis and a range of possible mortality rates on the other, reflecting high levels of uncertainty in these parameters. Since OCcAM was developed to be used with publicly available density data, OCcAM utilises mean peak seasonal *densities* rather than abundances as an input. The tool allows for a single, user-specified displacement rate and associated season-specific displacement mortality rates per species per parameter set (see supplementary information S1). These rates can readily be altered to derive predictions based on a range of possible input values.

### 3.3 Apportioning to reference population

The OCcAM tool has functionality to allow predicted mortalities to be put into the context of a focal population in line with methods currently applied in UK EIA. In some cases, total mortalities may be the required output. However, often there is a focal population of interest which may contribute only a subset of the birds expected to encounter and be impacted by the wind farm(s). In this case, mortalities must be apportioned to understand the relative impacts upon the reference population of interest.

The OCcAM tool includes three optional correction factors which can be applied to obtain appropriate final predicted mortalities for a reference population. These are: 1) the proportion of exposed birds that are expected to originate from the reference population; 2) the proportion of exposed birds that are expected to be adults (since seabird population counts are generally in adult birds); and, 3) the proportion of exposed adult birds expected to be sabbatical i.e. the proportion of adult birds not breeding in each year (since counts are often carried out at breeding sites during the breeding season when sabbatical birds may not be in attendance). Separate correction factors may be specified for each biologically relevant season, based on seasonal definitions presented by Furness (2015).

### 3.4 Tool design

The design and usage of the OCcAM tool is described in Supplementary Material S1.

## 4 Validation

Validated was carried out by comparing outputs of OCcAM analysis generated for Case Study 1 (Section 5), with predictions generated using the same input parameters independently with the respective standalone methods. Option 2 within the Band (2012) collision risk model spreadsheet was selected when generating the collision mortality estimates for validation purposes. All results were identical for displacement mortality and apportionment, while collision mortality estimates deviated slightly between the two approaches. This is due to the application of a user-defined function (UDF) within the Band (2012) spreadsheet which derives the proportion of birds at risk height from the flight height distributions defined within the spreadsheet. This functionality has not been included within OCcAM since it requires that the spreadsheet is macro-enabled, which can cause issues in terms of security, functionality and maintenance, and because inclusion of such features will lose some transparency for those users who are not familiar with scripting in Excel.

Validation results for the model runs for the Forth and Tay region in Case Study 1 are presented in Table 1.

**Table 1:** Comparison of collision risk predictions made using OCcAM versus the BTO Band spreadsheet.

| Region | Species | Parameter set | Birds at flight height OCcAM | Birds at flight height BTO | Collision estimate using OCcAM | Collision estimate using the BTO spreadsheet |
| --- | --- | --- | --- | --- | --- | --- |
| Forth and Tay | Kittiwake | PS1 | 0.1360 | 0.1314 | 70.88 | 69.49 |
| Forth and Tay | Kittiwake | PS2 | 0.0575 | 0.0563 | 157.16 | 154.73 |
| Forth and Tay | Gannet | PS1 | 0.1134 | 0.1093 | 48.04 | 47.34 |
| Forth and Tay | Gannet | PS2 | 0.0444 | 0.0434 | 97.25 | 96.17 |

## 5 Case study 1: Strategic level cumulative assessment for Scotland

### 5.1 Introduction

Where adverse effects on protected seabird populations are predicted, the Derogation Provisions under the Habitats Regulations provide a mechanism by which predicted impacts can be compensated. Since such compensation will generally be more effective at a regional rather than local scale, it is beneficial to be able to predict impacts over a large spatial scale. Compensatory measures can then be planned strategically based on the mortalities expected to be associated with the scale of buildout required to meet renewable energy targets. Such estimates would not be expected to represent exact numbers of mortalities, given uncertainties in future build-out scenarios and the many inherent conservatisms in the assessment process, but rather an indication of the order of magnitude of the number of birds that compensatory measures should be targeted towards.

Here we use OCcAM to generate a conservative prediction of the annual number of seabird mortalities that could result from the simultaneous operation of all existing and planned wind farm developments in Scottish waters.

### 5.2 Methods

Mortality rates from existing or planned Scottish offshore wind farms were predicted for the five OCcAM species, using the OCcAM tool. The tool was also used to apportion these mortalities to the adult breeding population belonging to Scottish SPAs.

Mortality was predicted separately for six regions of Scotland: Forth and Tay, Outer East, Moray Firth, North, Northeast and East, and West (see table 2). A separate OCcAM spreadsheet was set up for each region, and within each, separate parameter sets were used to predict (1) mortalities associated with operational wind farms, and (2) mortalities associated with wind farms that are planned, consented and under construction, based on status as defined in the RUK database (RenewableUK, 2024). Wind farms were separated this way since existing wind farms generally use older technology and therefore often have smaller turbines than more recent developments.

**Table 2:**
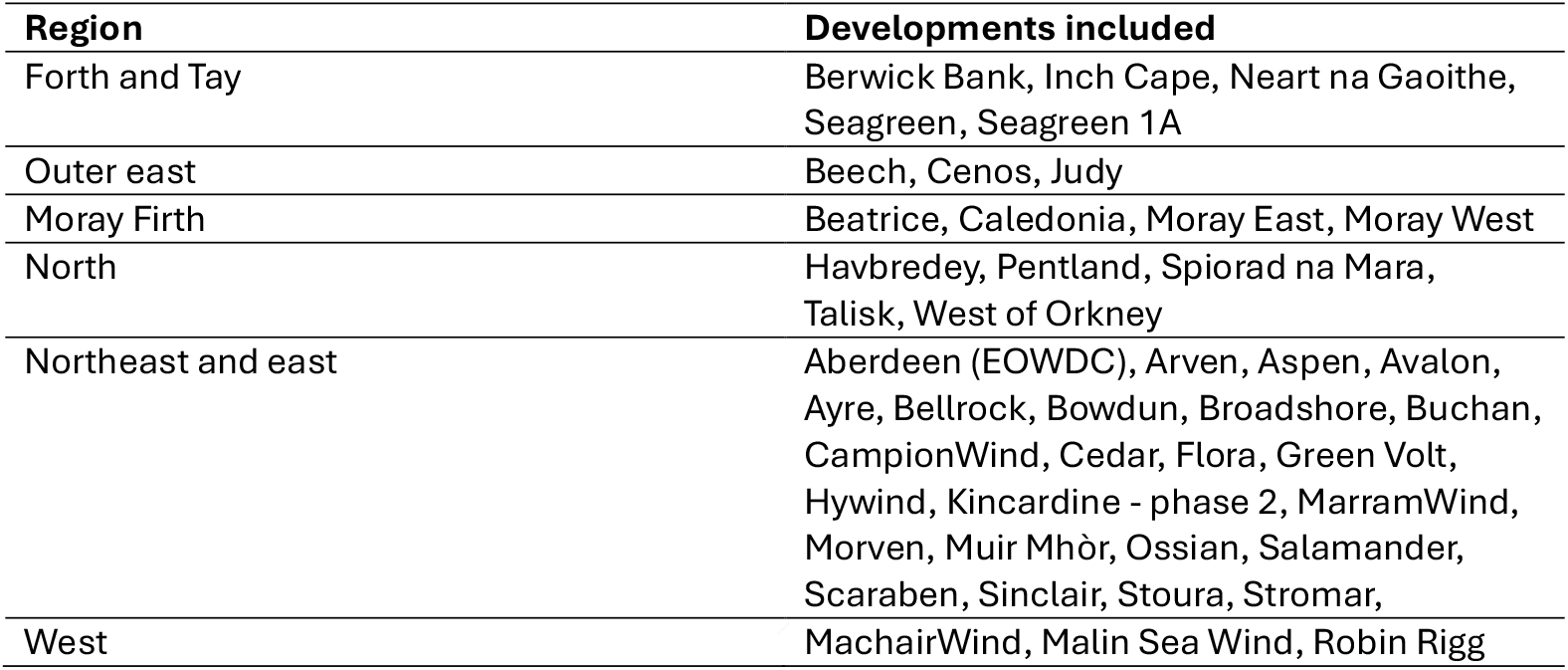
Regions used for OCcAM analysis and the developments included in each.

#### 5.2.1 Run and wind farm parameters

The latitude for each region was calculated as the centroid of the boundaries for those developments included within the region based on the 4C offshore database shapefile (TGS 4C Offshore, 2024). Area of windfarm footprints for each region-status combination was calculated as the sum of the array areas from the 4C database plus a 2km buffer. For existing wind farms, all user-defined wind farm collision inputs apart from air gap were taken from the RUK database (RenewableUK, 2024). Air gap was calculated as the reported tip height minus the rotor diameter from the RUK database. However, since air gap for CRM should be measured to mean sea level, and the datum to which tip height was measured is not stated within the RUK database, two metres were removed from the air gap to ensure that collisions were not underestimated due to the datum used. For wind farms in construction, consented or in planning, turbine numbers were taken from RUK if available or if not from 4C. Rotor diameter was taken from RUK where available, or assumed based on a current standard offshore turbine model, the Vestas V236-15. Air gap was assumed to be 30m based on offshore wind farm specifications from recently submitted EIAs.

#### 5.2.2 Bird impact parameters

Bird densities were calculated from publicly available density surfaces (Waggitt *et al*., 2020). For the collision risk modelling component within OCcAM, densities should represent birds in flight within the wind farm array areas. Monthly kittiwake and gannet densities were calculated for each region as the weighted average across the array areas of all of the wind farm polygons included in the region. Given that the densities provided by Waggitt *et al*., (2020) do not account for behaviour, these were then corrected to remove birds not in flight using correction factors derived from publicly available European Seabirds at Sea data (ESAS, 2023). ESAS records used to derive this ratio were based on all records falling within a convex hull enclosing the developments being considered. Where sample size of ESAS datapoints was too low to allow calculation of defensible ratios (N<100), global ratios calculated across all regions were applied. For bird densities required for displacement modelling, densities were determined from Waggit *et al*., (2020), as previously described for each array area plus a 2km buffer, and the month with the highest densities within each species-specific season was used as the input value. Other bird impact parameters were selected to mirror those currently recommended for use in EIA for offshore wind farms in Scotland (NatureScot, 2023b).

#### 5.2.3 Bird population parameters

Mortalities were compared to reference populations representing the adult breeding population at Scottish SPAs. Population sizes were taken from Burnell *et al*. (2023). For the breeding season, the proportion of fatalities expected to be attributable to this population was calculated as the proportion of SPA to non-SPA birds in Scotland, derived from Burnell *et al*. (2023). The proportion that was predicted to be adult was derived from a stable age structure calculated from population projection (Leslie) matrices constructed using demographic parameters presented in Horswill and Robinson (2015). For the non-breeding season, data provided in Furness (2015) was used to derive proportions of mortalities attributable both to the Scottish SPAs and to the adult portion of the population. Sabbatical corrections were applied for both breeding and non-breeding seasons based on those used in recent Scottish offshore wind EIAs (Green Volt, 2023).

All OCcAM spreadsheets used for this analysis including input parameters are provided in Supplementary Material S2.

### 5.3 Results

For kittiwake, OCcAM indicates that approximately 386 mortalities of breeding adults at Scottish SPA populations could be associated with offshore windfarm development annually, representing 0.19% of the current SPA breeding adult population.

Approximately 483 annual gannet SPA breeding adult mortalities are predicted representing 0.10% of the current SPA breeding adult population (Figure 1).

**Figure 1:**
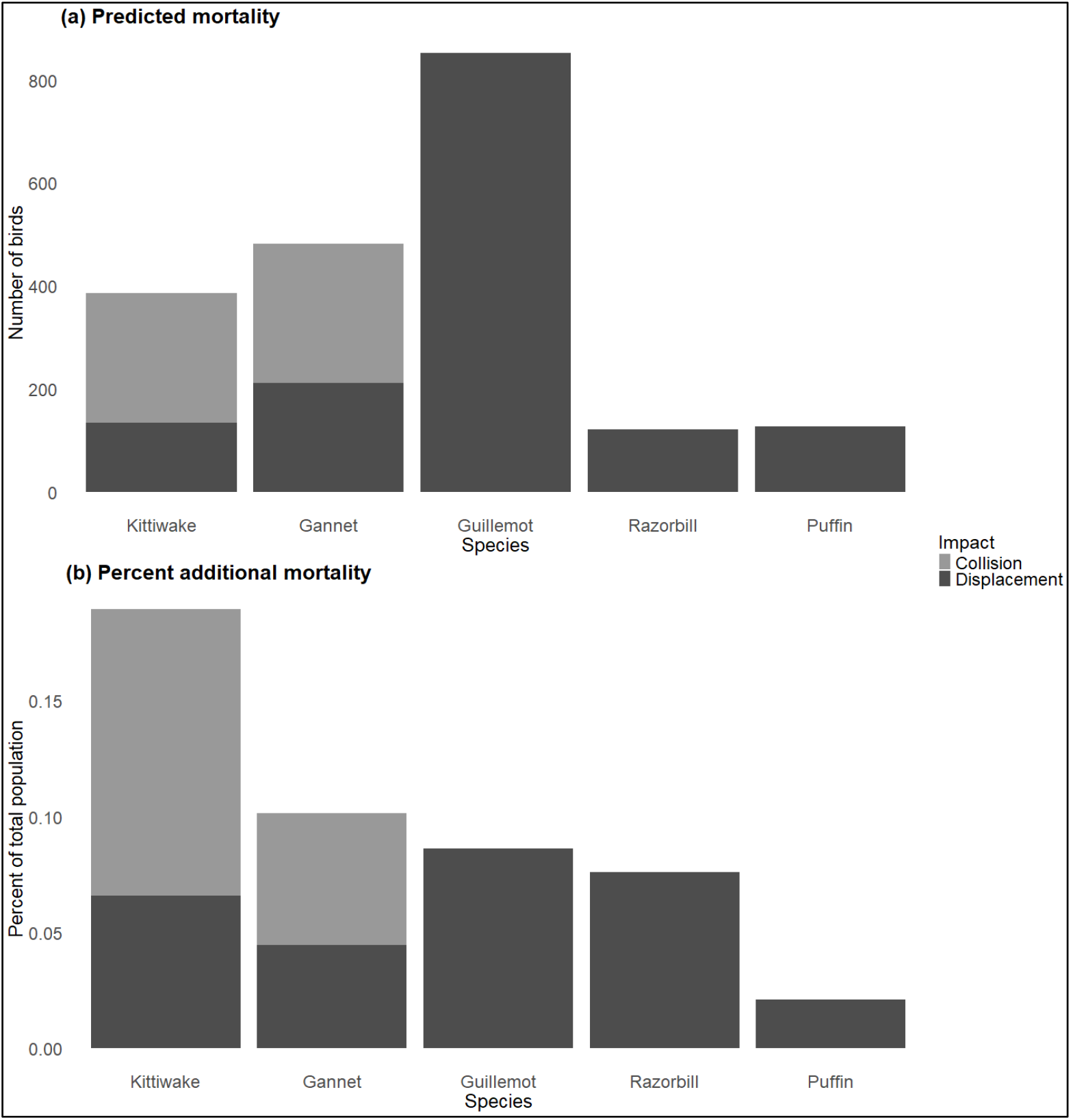
The number of mortalities predicted to be attributable to (a) the Scottish breeding adult SPA population and (b) the percentage of the total population size represented

For guillemot, razorbill and puffin, 852, 122 and 128 SPA breeding adult mortalities are predicted respectively from distributional responses, representing 0.09%, 0.08% and 0.02% of the Scottish SPA breeding adult population size respectively (Figure 1).

### 5.4 Conclusions

Case study 1 provides an indicative number of potential fatalities associated with the build out of all existing and potential upcoming projects in Scottish waters, both overall, and related specifically to birds associated with protected populations. However, the combined capacity of the developments included in the analysis sums to 45.56GW which far exceeds any targets set by the Scottish Government so far. It is extremely unlikely that all of this capacity will be realised nor that all of the developments will be operational at the same time. As such this analysis represents a conservative assessment. The inclusion of operational sites is an additional conservatism given that any annual mortalities attributable to these sites will contribute to what is currently considered as baseline fatality rates for these species in the area.

That said, a number of assumptions have been made to derive the input parameters, which may lead to under- or over-estimation of impacts. One key source of uncertainty is the density estimates used in the modelling. The dataset used exploits aerial and boat-based survey data collected for other purposes, between 1980 and 2018. The surfaces generated may therefore not be reflective of current patterns of seabird densities and may also be subject to biases from unbalanced spatial and temporal survey coverage. However, at this time, the Waggitt *et al*., (2020) density surfaces are the only published data covering UK waters and are based on the most extensive datasets currently available. As such, they have also recently been utilised for the purposes of strategic planning by the UK Crown Estates (NIRAS, 2024). Use of publicly available ESAS data to calculate proportion of birds in flight will introduce further uncertainty due to potential biases associated with patchy coverage and the spatial scales over which these corrections were calculated. Ongoing studies funded by the Offshore Wind Evidence and Change (OWEC) programme are expected to generate more temporally relevant predictions that could easily be used to generate updated density estimates for use with OCcAM (OWEC, 2024).

Despite these factors, the numbers generated by OCcAM can be used as an indicative target to inform the extent of compensatory measures that should be put in place to offset any negative impacts of offshore wind farm development over coming years in Scottish waters. In this case, the analysis suggests that compensatory measures should target four to five hundred kittiwake and gannet per year, less than a thousand guillemot per year and one to two hundred razorbill and puffin per year to offset mortality associated with the development of offshore windfarms. The results can be updated as new data (for example seabird density maps) or parameter updates (for example updated displacement, displacement mortality or avoidance rates) become available, allowing targets to be readily reviewed and updated in light of new information.

## 6 Case study 2: Comparing as-built versus consented layout

### 6.1 Introduction

EIA for offshore wind farm applications generally happens in advance of the determination of the final project design, and is therefore based on indicative parameters, often representing a design “envelope” within which the final buildout parameters are expected to fall. Consenting decisions are made considering the worst-case scenario presented within the envelope. These publicly available mortality estimates are then generally taken forward for use in cumulative assessment (OWEC, 2021). However, differences in turbine number, size and height between the worst-case consented scenario and the final built-out scenario can make a big difference to predictions of mortality arising from collision, and changes in the array area footprint will affect the number of birds predicted to be displaced. Use of impacts predicted during EIA for consent can therefore result in predictions of greater annual mortality from existing wind farms than would be expected based on built-out parameters. This could potentially lead to reduced scope for consenting of further wind energy development as a result of being unable to conclude no adverse effects on site integrity for protected seabird populations. In order to meet renewable energy generation targets, resolving this issue of so-called “headroom”, i.e. the difference in expected mortalities associated with consented parameters versus those that would be predicted based on the as-built wind farm design, has become a key focus area in the UK (OWEC, 2021).

Here we present a case study in which we use OCcAM to compare cumulative impacts calculated based on consented and as-built parameters, demonstrating how OCcAM can be applied to update mortality estimates based on built out parameters and providing an indication of the magnitude by which use of consented versus built out parameters might influence predictions of cumulative effects.

### 6.2 Methods

OCcAM was used to predict cumulative collision and displacement impacts upon seabirds from currently operational wind farms in the Moray Firth region of Scotland based on (1) the worst-case scenario consented parameters (representing those generally used in project-specific cumulative impact assessments) and (2) the final built out parameters. Three offshore windfarms within the Moray Firth regional grouping are currently operational; Moray East, Moray West and Beatrice.

Many user-defined and default parameters can be edited within the OCcAM tool. However, this case study focuses on those wind farm parameters that are known to be sensitive in estimation of bird impacts, and which can differ dramatically between the worst-case consented design and the final buildout. These parameters are: (1) turbine number, (2) rotor diameter, (3) air gap and (4) array area plus 2km buffer.

Consented and as-built parameters for the three wind farms were provided by RUK. Once parameters had been extracted, overall values for the region were calculated by summing turbine numbers and array areas across developments in each scenario, and by taking weighted averages of the rotor diameter and air gap for each scenario. Resulting input parameters are presented in Table 2.

**Table 2:** Worst-case consented and as-built input wind farm parameters calculated for the Moray Firth region of Scotland.

| Scenario | Number of turbines | Rotor diameter | Air gap* | Area of wind farm footprint + 2km buffer |
| --- | --- | --- | --- | --- |
| Worst-case consented | 548 | 142.8 | 24.4 | 1087.128 |
| As built | 244 | 174.8 | 32.6 | 1047.386 |

\* Note that air gaps in this study differ from those in Case Study 1 since the resolution of this analysis is finer, so more detailed investigation into precise parameters was applied.

OCcAM was then run using identical parameters to those in Case Study 1 for the Moray Firth region, but with the relevent updated wind farm specifications for each scenario.

The OCcAM spreadsheet used including input parameters are provided in Supplementary Material S2.

### 6.3 Results

For all species, predicted mortalities were lower when derived from the as-built parameters compared to those derived from the worst-case consented parameters (Figure 2). Differences in predictions for guillemot, razorbill and puffin, considered to be at risk from displacement mortality only, are small, with predictions based on consented values being 3.8% higher than those predicted based on the as-built scenario, which translated to a maximum difference in total mortalities of almost 6 individuals, for guillemot (Figure 2).

**Figure 2:**
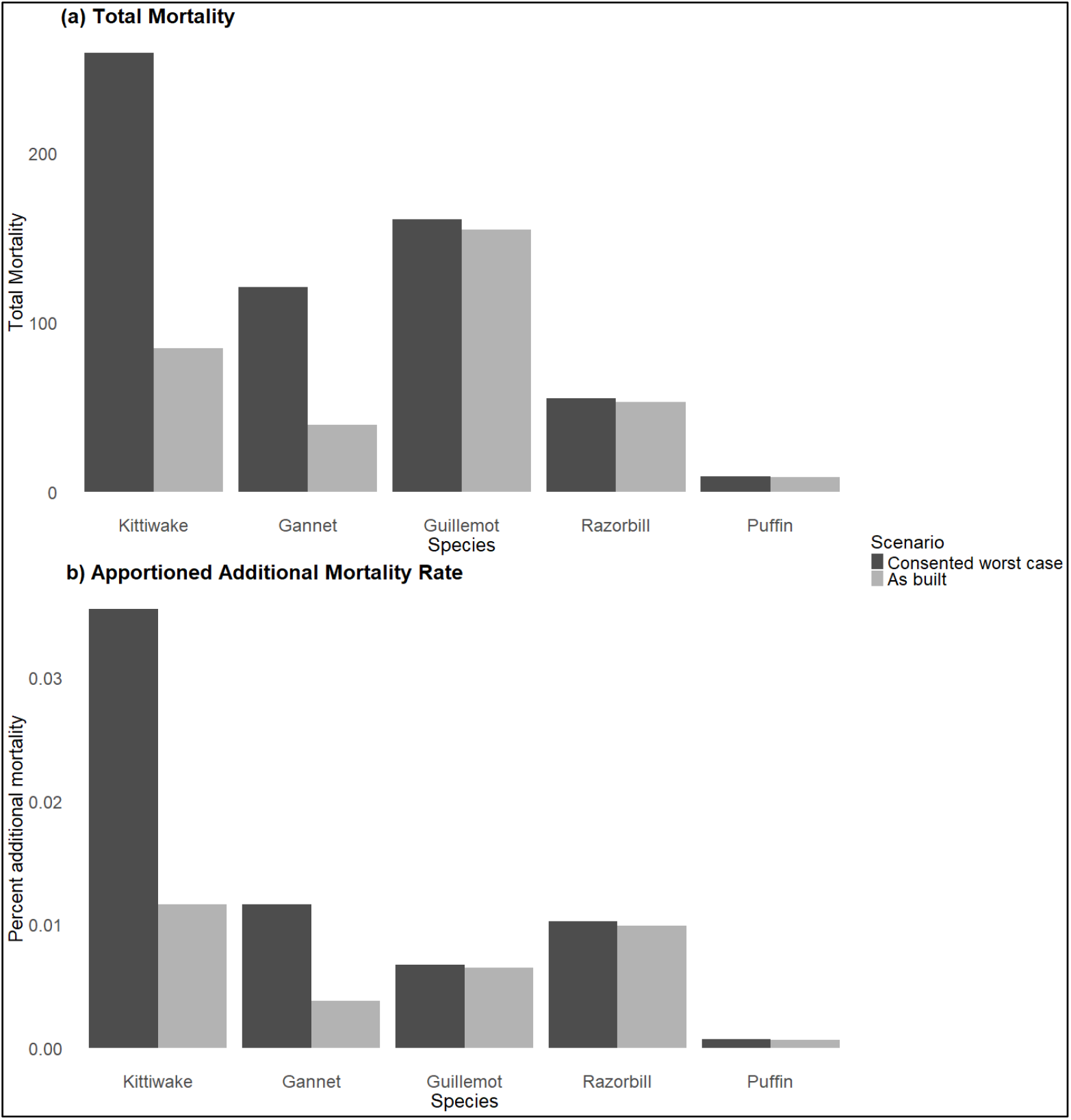
Cumulative total mortality (a) and apportioned percent additional mortality (b) predicted for the Moray Firth region of Scotland based on consented worst-case scenario design parameters and as-built design parameters.

For kittiwake and gannet, the differences are considerably larger, with more than 200% higher mortality predicted based on the consented values compared to as built, translating to absolute difference differences in total mortalities of 174 and 81 individuals respectively. This results in percent additional mortality from consented design parameters on the Scottish SPA breeding adult population that is around 205% higher using the consented as opposed to the as-built design parameters.

### 6.4 Conclusions

There are often substantial differences between the design parameters included in project-level impact assessments used for consenting purposes, and those associated with the realised build-out scenario for the same project. Parameters subject to change include turbine number, size and height, as well as array area footprint, all of which are sensitive parameters in determining predicted seabird impacts.

In this example, the footprint of the array area remained the same for two out of the three developments included in the analysis, and the area of the third reduced by a very small percentage (Table 2). Since this is the only parameter of the four considered used in estimation of mortality arising from distributional responses, estimated mortality for the species for which only distributional responses are considered, i.e. guillemot, razorbill and puffin, changed very little between the worst-case consented and the realised as-built scenario.

However, differences in parameters used within collision risk modelling were much greater, and updating these parameters for kittiwake and gannet dramatically reduced the predicted mortality rates for this region.

These results demonstrate that use of mortality estimates presented in project-level assessments used for consent are not appropriate for use in cumulative assessment and may lead to substantial overestimates of mortality, limiting the capacity to consent further projects. OCcAM provides a quick and easy method to update existing assessments, providing a means to generate impact predictions based on as-built parameters using a methodological approach that will give comparable results to that used during the EIA process (where the matrix method and the Band collision risk model have been applied).

## 7 Case study 3: Flexible scoping of a project

### 7.1 Introduction

During the development of an offshore wind farm site, many decisions are made relating to the profile of the project itself. These decisions primarily include the siting, numbers, size and height of turbines, including air gaps between the rotor and the sea, all of which may have a large impact on numbers of bird mortalities predicted. OCcAM provides a method to use available bird density data to investigate the impacts of such decisions on the potential to adversely impact upon seabird populations by allowing rapid modelling of a number of potential scenarios. These models can then be updated during the development of the project and alongside the formal EIA process, including site-specific datasets as they become available. In addition to helping reduce potential impacts by embedding mitigation by design from the earliest stages, outputs can also provide useful evidence to inform early engagement and consultation with regulators and stakeholders to reduce consenting timeframes post-application.

Here we present a case study in which a hypothetical wind farm off the Scottish coast is designed to minimise impacts on seabirds based on outputs generated using OCcAM.

### 7.2 Methods

A scenario was simulated in which the developer of a hypothetical wind farm wanted to understand the broad-scale magnitude of (1) changing the shape of the array area, (2) using different turbine models to achieve the desired energy capacity for the site, and (3) scenarios using different air gaps.

Two 347km^2^ array areas were generated, both based around a centroid of 57°52’58.008’’ N, 1°23’2.824 W. One array area consisted of a square with sides of 18.6km, whilst the other consisted of an upright rectangle of 40km x 8.65km. Buffering these areas by 2km gave displacement areas of 507.2km^2^ for the square array area and 553.2km^2^ for the rectangular array area.

OCcAM runs were carried out for 18 scenarios, representing all possible combinations of (1) two array area shapes, (2) three turbine specifications and (3) three different air gaps. This was achieved using six OCcAM spreadsheets.

Turbine specifications were based on three common turbine models in use offshore: the Vestas V236-15 turbine, with a capacity of 15MW per turbine and a rotor diameter of 236m, the GE Haliade-X 14MW turbine, with a capacity of 14MW per turbine and a rotor diameter of 220m and the Siemans Gamesa SG 11.0-193 DD turbine, with a per turbine capacity of 11MW and a rotor diameter of 193m. The number of turbines of each model included in each scenario was calculated by dividing an assumed target total site capacity of 1500 MW by the capacity of the individual turbines for that model. This gave scenarios of 100 x 15MW Vestas turbines, 107 x 14MW GE turbines and 136 x 11MW Siemens turbines.

The three air gaps used were 30m, 35m and 40m above mean sea level which are representative of air gaps for currently proposed and recently constructed offshore wind farms.

Since the hypothetical array areas fall within the Scotland – North East and East region of Scottish waters used in Case Study 1, all bird input parameters replicated those in the analysis carried out for this region under Case Study 1. Since broad-scale bird densities were used, no account was made for potential changes in bird densities based on the two different array area shapes. OCcAM spreadsheets including input parameters are provided in Supplementary Material S2.

### 7.3 Results

The results of the analysis demonstrate that in this case, the developer could minimise their impact to all species by choosing the more compact square-shaped array area, building fewer, larger turbines and maximising the air gap (Figure 3). For the auk species, only the shape of the array area is important, since this is the only parameter that feeds into displacement estimates. However, by having a square rather than a rectangular array area, the predicted impact on guillemot reduced by 8%. For gannet and kittiwake, for which OCcAM has been used to predict both displacement and collision in this case, air gap is by far the most important determinant of overall impacts – with an increase in air gap from 30m to 40m resulting in approximately 40% and 37% reductions in overall mortality for kittiwake and gannet respectively.

**Figure 3:**
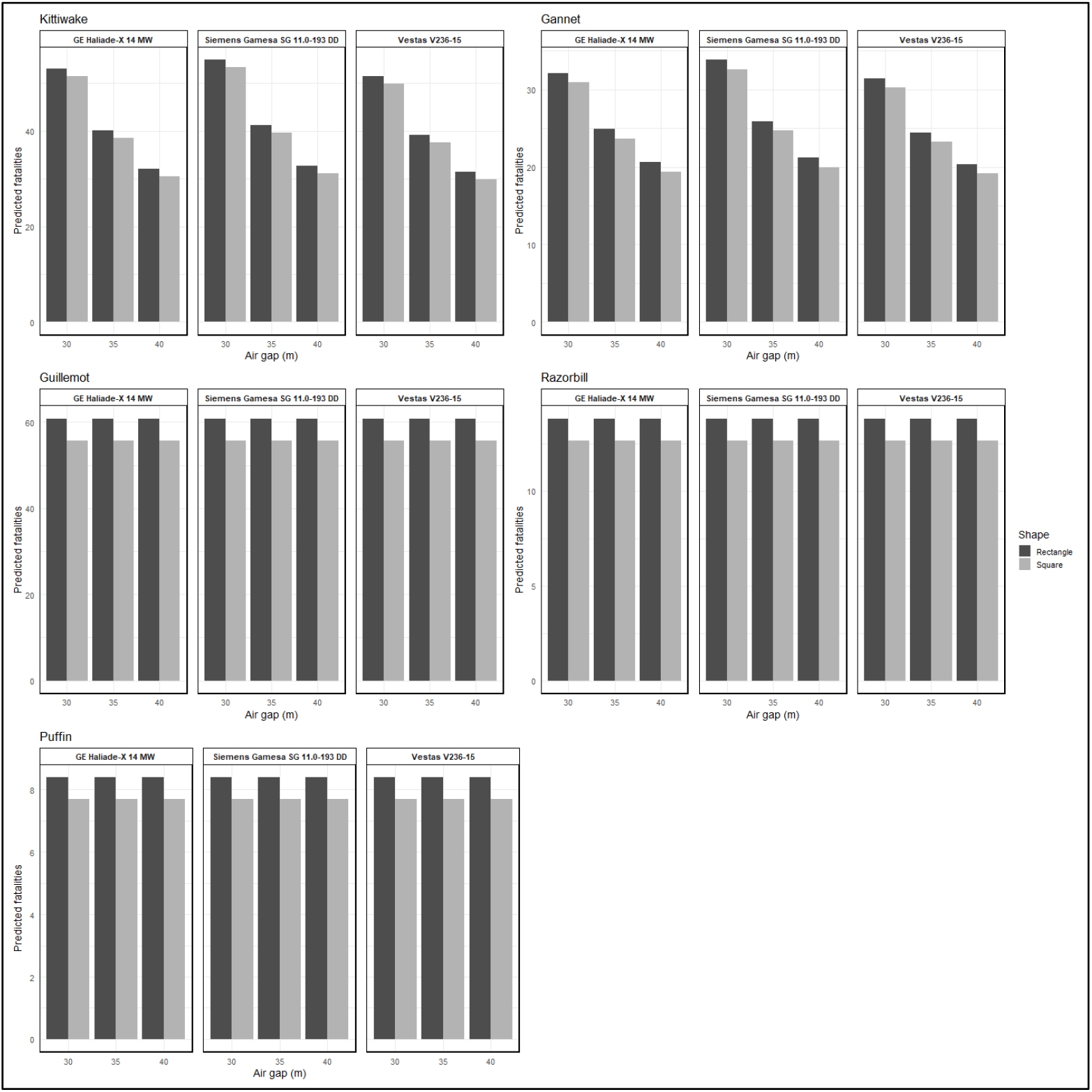
Difference in predicted fatalities for the five species represented within OCcAM under different wind farm configurations.

### 7.4 Conclusions

OCcAM provides a practical alternative to using existing assessment methods to indicate the implications of different design choices on predicted mortality. The format of the tool allows simultaneous and instantaneous comparison of multiple designs to both collision and displacement impacts to five species, providing 30 different informative analyses in a single run. The outcomes of these analyses can then be included alongside other considerations and constraints feeding into the design process.

## 8 Discussion

### 8.1 Utility and tool applications

OCcAM leverages two industry-standard methods applied within the UK to rapidly assess the potential impacts of offshore windfarms on seabirds within a simple and user-friendly tool.

OCcAM has not been designed to replicate the more complex aspects of analysis currently recommended by UK regulators for project-level EIA (NatureScot, 2024; SNCBs, 2024, 2022), and the inclusion of stochasticity has been intentionally omitted in order to ensure the simplicity, efficiency and flexibility of the tool. The tool aims to allow technical and non-technical users alike to quickly and easily generate robust broad-scale predictions, as well as to facilitate a deeper understanding of the underlying drivers of variability in those predictions and the potential implications of misspecification of parameters in the context of overall mortality rates. If OCcAM were to be used to generate predictions for a specific project using the same parameters as were used for project-only impact assessment using the matrix displacement approach and the sCRM (Caneco *et al*., 2022), it would be expected to give very similar results, with differences in predictions deriving only from the ways in which stochasticity associated with input parameters are incorporated into the sCRM model.

Population modelling, often carried out as the final stage of the EIA in order to understand long-term population-level effects, has not been incorporated into the tool due to the increased number of parameters and level of complexity that this would involve. However, the “percentage of population impacted” output represents the percentage point change to the adult annual survival rate recommended for use by NatureScot as a threshold to determine species for which population modelling should be carried out (set to 0.02% at the time of writing (NatureScot, 2023c)), and is also the proportion adult mortality rate used in population modelling. OCcAM can therefore be used to screen populations for which population modelling should be carried out, as well as providing inputs that can then be directly taken forward into population modelling.

It is expected that the recommended input parameters for the methods applied within OCcAM will change over time and OCcAM has been future proofed with this in mind. Many of the most sensitive parameters, and therefore those which may be most likely to be updated, have been incorporated as user-defined parameters. These include avoidance rates, displacement rates and displacement mortality rates. In addition, default parameters are easily viewed and, whilst “protected” to prevent accidental edits, can easily be “unprotected” (Review tab in Excel) and updated as well.

The original purpose of OCcAM was to predict the magnitude of potential impacts of offshore wind development at a regional scale. As such, it can be used to support planning and decision-making at the strategic level by facilitating a high-level understanding of potential cumulative impacts of current, planned and future developments over a large area where suitable input data are available, as demonstrated in Case Study 1. This use case would be beneficial for marine planning, for example by (1) potentially supporting decisions such as how many, what size and where new developments may be built out to minimise impacts to seabirds; and (2) predicting the magnitude of compensatory measures that may be required under the Habitats Regulations to offset impacts of existing and planned offshore windfarm development, allowing appropriate allocation and effective management of marine recovery funds to support protected seabird populations.

Where cumulative impact assessment is based on extracting predicted impacts from previous EIAs, as is often the case for project-specific cumulative EIA, OCcAM can also be a useful tool to quickly understand differences between estimates based on the worst-case scenarios represented within project design envelopes and the as-built scenarios reflecting the realised situation, as in Case Study 2. This allows for more accurate and proportionate cumulative impact assessment, potentially removing conclusions of adverse effects on site integrity and therefore the need to deliver unwarranted compensation measures.

As demonstrated in Case Study 3, OCcAM can be used as a tool by developers to embed ornithological considerations into the design of an offshore wind farm from the very start of project planning by highlighting the key risks for the species included and demonstrating how these can change as a result of altering design parameters. An early indication of potential impacts from OCcAM will also facilitate early consultation with regulators and other stakeholders on the key expected impacts of a proposed project, providing a data-driven basis for discussion of requirements for EIA. During project development, general density data can be replaced with site-specific data to refine predictions as a more detailed understanding of the situation at the site emerges. This allows project engineering parameters to evolve alongside an understanding of what the key risks are from an ornithological perspective.

Use of OCcAM for the cases highlighted above could result in marked reductions in timescales and costs for ornithological impact assessment. A shorter planning and consenting period for offshore wind farm developments can lead to significant societal, environmental and industry benefits through a quicker move to renewable energy sources. DEFRA has recently estimated that reducing consenting timeframes from the current scenario of 5 to 7.5 years in the UK by 1-2 years would result in a £2.4 billion reduction to society from carbon savings(DEFRA, 2025b).

Case studies presented here have focused on usage within the UK, where the offshore renewable energy industry is mature and guidance and industry-standards for ornithological impact assessment are well-established. However, OCcAM also provides a tool that can be taken up or adapted for other regions where the offshore wind industry is less mature, and where standard practices for ornithological impact assessment have yet to be established. OCcAM can expedite uptake of these established models, for example in emerging markets, in a way that is quickly implemented and easily interpretable to stakeholders, leading to efficient development of appropriate market-specific processes.

### 8.2 Possible extensions and data requirements towards wider tool applicability

OCcAM currently includes the option to predict mortality for five seabird species, selected due to their relevance across the UK and northern Europe, with a particular focus on Scotland (McGregor *et al*., 2022; Scottish Government, 2024). OCcAM could readily be extended to include additional focal species, or separate versions could be developed to cover key species for different regions or concerns. In some instances, it would also be possible for a user to independently replace the species within OCcAM for other species for which the same number of seasons were required, by updating data in the default data tab.

In the case studies presented here, publicly available density surface data covering the area of interest have been used to derive densities for use within the tool. Model outputs are highly sensitive to input densities, so it is important that users understand the potential limitations associated with densities used and apply this knowledge to ensure appropriate inference from outputs. Increased knowledge of distributions of seabirds at sea is a priority research area in many countries, and OCcAM can readily be updated with new densities as these become available.

From a UK perspective, the OWEC Planning Offshore Wind Strategic Environmental Impact Decisions (POSEIDON) project is expected to release new density models for key receptors in the short-term based on current and systematically collected data (OWEC, 2024), and these will supersede the Waggitt *et al*., (2020) densities applied here with more robust results for several species.

For other regions and use cases, there are a number of studies presenting regional and EEZ-wide density estimates which could be used to generate predictions, including those derived from the ObSERVE programme in Ireland (Jessop *et al*., 2018; Paradell *et al*., 2024), those derived as part of the Masterplan Ecologische Monitoring Lange Termijn Wind op Zee in the Netherlands (Poot *et al*., 2011) and densities estimated by Pettex *et al*. (2017) from aerial survey campaigns in the Bay of Biscay and the English Channel.

OCcAM was developed to facilitate rapid, large-scale prediction of mortalities, giving an indication of the magnitude of potential impacts. There are therefore several default parameters, such as the default turbine specifications linked to the user-defined rotor diameter, which have been selected based on reducing the amount of effort required to extract appropriate parameters when applying the tool. However, depending on the application, these parameters may be readily available. Whilst it is straightforward to overwrite default parameters within OCcAM, depending on uptake and usage of the tool, it may be beneficial to build in an option to select between default and user-defined options for these parameters to avoid updating the default parameters tab becoming a standard component of the usage of OCcAM.

## 9 Conclusions

Quantifying and minimising likely impacts of proposed offshore wind farms on seabirds in a timely and transparent way is crucial for ensuring compliance with environmental legislation and safeguarding seabird populations, whilst simultaneously facilitating wind energy development to support climate goals. OCcAM provides a method to quickly generate mortality estimates, with numerous potential applications in the UK and elsewhere, from broad-scale predictions that can support strategic-level decision making, to facilitate consenting, from site feasibility through to condition discharge via rapid scoping of changes to wind farm configurations and assessment parameters. As a publicly available tool developed from within the industry with the end user in mind, OCcAM is available and readily implementable for all stakeholders within the renewable energy sector.

## Notes

### Competing Interest Statement

The authors have declared no competing interest.

### Funding

- SSE Renewables

## References

Band, B., 2012. Using a collision risk model to assess bird collision risks for offshore windfarms. Report by British Trust for Ornithology (BTO).

Bolton, P., 2025. Clean power targets: UK House of Commons research briefing. UK House of Commons.

Bradbury, G., Trinder, M., Furness, B., Banks, A.N., Caldow, R.W.G., Hume, D., 2014. Mapping seabird sensitivity to offshore wind farms. PLOS ONE. 10.1371/journal.pone.0106366

BTO, 2013. SOSS Collision Risk Modelling Excel Spreadsheet [Excel file].

Burnell, D., Perkins, A.J., Newton, S.F., Bolton, M., Tierney, T.D., Dunn, T.E., 2023. Seabirds Count: a census of breeding seabirds in Britain and Ireland (2015– 2021). Lynx Nature Books, Barcelona.

Caneco, B., Humphries, G., Cook, A.S.C.P., Masden, E.A., 2022. Estimating bird collisions at offshore windfarms with stochLAB.

COP28, IRENA, GRA, 2023. Tripling renewable power and doubling energy efficiency by 2030: Crucial steps towards 1.5°C, International Renewable Energy Agency, Abu Dhabi.

Dalal-Clayton, B., Scott-Brown, M., 2024. Improving decision-making for the energy transition: Guidance for using Strategic Environmental Assessment. International Association for Impact Assessment.

DEFRA, 2025a. Strategic compensation measures for offshore wind activities: Marine Recovery Fund interim guidance.

DEFRA, 2025b. Marine Recovery Fund Options Assessment. DEFRA.

Drewitt, A.L., Langston, R.H.W., 2006. Assessing the impacts of wind farms on birds. Ibis 148, 29–42. 10.1111/j.1474-919X.2006.00516.x

ESAS, 2023. European Seabirds At Sea dataset [dataset]. ICES, Copenhagen, Denmark.

EU, 2023. Directive (EU) 2023/2413 of the European Parliament and of the Council of 18 October 2023 amending Directive (EU) 2018/2001, Regulation (EU) 2018/1999 and Directive 98/70/EC as regards the promotion of energy from renewable sources, and repealing Council Directive (EU) 2015/652. Off. J. Eur. Union.

Furness, R.W., 2015. Non-breeding season populations of seabirds in UK waters; Population sizes for Biologically Defined Minimum Population Scales (BDMPS). (Natural England Commissioned Reports No. Number 164).

Galparsoro, I., Menchaca, I., Garmendia, J.M., Borja, Á., Maldonado, A.D., Iglesias, G., Bald, J., 2022. Reviewing the ecological impacts of offshore wind farms. Npj Ocean Sustain. 1, 1. 10.1038/s44183-022-00003-5

Green Volt, 2023. Green Volt Offshore Windfarm: Report to Inform Appropriate Assessment.

Horswill, C., Robinson, R.A., 2015. Review of seabird demographic rates and density dependence. (JNCC Report No. No: 552). Joint Nature Conservation Committee, Peterborough.

Jessop, M., Mackey, M., Luck, C., Critchley, E., Bennison, A., Rogan, E., 2018. The seasonal distribution and abundance of seabirds in the western Irish Sea. Department of Communications, Climate Action and Environment, and National Parks & Wildlife Service, Department of Culture, Heritage & the Gaeltacht, Ireland.

McGregor, R., Trinder, M., Goodship, N., 2022. Assessment of compensatory measures for impacts of offshore windfarms on seabirds. A report for Natural England. Natural England Commissioned Reports. (No. NECR431).

Morgan, R.K., 2012. Environmental impact assessment: the state of the art. Impact Assess. Proj. Apprais. 30, 5–14. 10.1080/14615517.2012.661557

NatureScot, 2024. Advice on marine renewables development.

NatureScot, 2023a. Guidance Note 8: Guidance to support Offshore Wind Applications: Marine Ornithology Advice for assessing the distributional responses, displacement and barrier effects of Marine birds.

NatureScot, 2023b. Guidance Note 7: Guidance to support Offshore Wind Applications: Marine Ornithology - Advice for assessing collision risk of marine birds.

NatureScot, 2023c. Guidance Note 11: Guidance to support Offshore Wind Applications: Marine Ornithology - Recommendations for Seabird Population Viability Analysis (PVA).

NIRAS, 2024. Round 5 Plan HRA RIAA: Report to inform Appropriate Assessment: CF350 Birds detailed assessment: CF357 Seabird densities.

O’Brien, S., 2025. Streamlining Ornithology Impact Assessments: What changes would be beneficial?

OWEC, 2024. The Offshore Wind Evidence and Change Programme Annual Report 2024.

Paradell, G., Rogan, O., Jessopp, M., 2024. The seasonal distribution and abundance of seabirds, cetaceans and other megafauna off the south and southwest Irish coast (No. Department of the Environment, Climate and Communications and Department of Housing, Local Government and Heritage, Ireland. 133pp).

Parker, J., Fawcett, A., Banks, A., Rowson, T., Allen, S., Rowell, H., Harwood, A., Ludgate, C., Humphrey, O., Axelsson, M., Baker, A., Copley, V., Robertson, A., Hodgkiss, R., Berridge, R., Farmer, R., 2025. Offshore Wind Marine Environmental Assessments: Best Practice Advice for Evidence and Data Standards. Phase III: Expectations for data analysis and presentation at examination for offshore wind applications. Natural England. Version 2.

Pettex, E., Laran, S., Authier, M., Blanck, A., Dorémus, G., Falchetto, H., Lambert, C., Monestiez, P., Stéfan, E., Van Canneyt, O., Ridoux, V., 2017. Using large scale surveys to investigate seasonal variations in seabird distribution and abundance. Part II: The Bay of Biscay and the English Channel. Deep Sea Res. Part II Top. Stud. Oceanogr. 141, 86–101. 10.1016/j.dsr2.2016.11.012

Poot, M.J.M., Fijn, R.C., Jonkvorst, R.J., Heunks, C., Collier, M.P., de Jong, J., van Horssen, P.W., 2011. Aerial surveys of seabirds in the Dutch North Sea May 2010 – April 2011 Seabird distribution in relation to future offshore wind farms. (No. 10–235). Bureau Waardenburg bv.

Ramirez, L., Fraile, D., Brindley, G., 2020. Offshore Wind in Europe: Key trends and statistics 2019.

RenewableUK, 2024. [dataset] https://www.renewableuk.com/energypulse/database/ Accessed: 15/11/2024

Scottish Government, 2024. The Scottish Seabird Vulnerability Report.

SNCBs, 2024. Joint advice note from the Statutory Nature Conservation Bodies (SNCBs) regarding bird collision risk modelling for offshore wind developments. JNCC, Peterborough.

SNCBs, Statutory Nature Conservation Bodies, 2022. Joint SNCB Interim Displacement Advice.

Tapia-Harris, C., Evans, T., 2024. Feasibility of strategic ornithological compensatory measures in the Scottish context.

TGS 4C Offshore, 2024. [dataset] https://map.4coffshore.com/offshorewind/ Accessed: 08/11/2024.

Waggitt, J.J., Evans, P.G.H., Andrade, J., Banks, A.N., Boisseau, O., Bolton, M., Bradbury, G., Brereton, T., Camphuysen, C.J., Durinck, J., Felce, T., Fijn, R.C., Garcia-Baron, I., Garthe, S., Geelhoed, S.C.V., Gilles, A., Goodall, M., Haelters, J., Hamilton, S., Hartny-Mills, L., Hodgins, N., James, K., Jessopp, M., Kavanagh, A.S., Leopold, M., Lohrengel, K., Louzao, M., Markones, N., Martínez-Cedeira, J. Ó, Cadhla, O., Perry, S.L., Pierce, G.J., Ridoux, V., Robinson, K.P., Santos, M.B., Saavedra, C., Skov, H., Stienen, E.W.M., Sveegaard, S., Thompson, P., Vanermen, N., Wall, D., Webb, A., Wilson, J., Wanless, S., Hiddink, J.G., 2020. Distribution maps of cetacean and seabird populations in the North-East Atlantic. J. Appl. Ecol. 57, 253–269. 10.1111/1365-2664.13525

