## Supplementary information S1 for "OCcAM: A tool for rapidly assessing impacts of offshore wind farms on seabirds"

### Supplementary Information 1: OCcAM Tool design

The OCcAM tool allows analysis of up to three different parameter sets in a single run. Parameter sets can be used either to investigate three different realisations of the same scenario e.g. three different assumed wind farm configurations, or to generate a single mortality estimate based on sub-strata, for example, wind farms in areas of high, medium and low bird density.

The OCcAM tool comprises an introduction tab and eleven analysis tabs which can be categorised into four categories: those that take in user inputs, a tab which displays default parameters, tabs which carry out intermediate calculation steps, a results tab which presents the final predictions and a run log tab. These tabs are described in more detail below.

#### 1 User input tabs

OCcAM includes three tabs within which the user must enter input parameters required for the runs. The *run and wind farm* tab includes parameters relating to the setup of the tool for the run and to the site and wind farm itself. The inputs entered here are used for both collision risk modelling and displacement analysis. The *bird impacts* tab requires input of bird densities and behavioural parameters as well as displacement mortality rates, also feeding into both collision and displacement predictions. The *bird populations* tab is only used where comparisons should be made to a reference population and requires the size of the reference population and the correction factors to be used. These inputs are used to apportion predicted mortalities to the reference population. All user input parameters and their explanations are presented in Table 1.

*Table 1: OCcAM user-defined parameters*

| Parameter | Unit | Description |
|---|---|---|
| **Run and wind farm parameters** | | |
| Latitude of region centroid | Decimal degrees | The latitude of the centroid of the region of interest, used to calculate daylight hours for collision risk modelling. |
| Number of turbines | Turbines | The total number of turbines. |
| Rotor diameter | Metres | The diameter of the turbines being considered - may be an average where multiple turbine specifications are to be included in the analysis. |
| Air gap | Metres | The distance between the bottom of the rotor-swept volume and the sea surface relative to mean sea level. This may be an average where multiple turbine specifications are included in the analysis. |
| Total area of wind farm footprint | Square kilometres | The area within which birds are expected to display distributional responses to the wind farms under consideration. This is often calculated as the array area plus a 2 km buffer for each wind farm considered, depending upon the species concerned. |
| **Bird impacts - collision** | | |
| Include macro-avoidance for gannet | Yes/No | If this option is selected then the gannet collision estimate will be reduced by the proportion expected to be displacement from the array area according to the displacement input parameters. |
| Kittiwake and gannet monthly in-flight densities | Birds per square kilometre | Estimated densities of birds in flight within the array area, used for collision modelling. |
| Kittiwake and gannet avoidance rate | Proportion | The proportion of birds flying within the array area expected to take avoidance action to avoid a collision that otherwise would have taken place. |
| Kittiwake and gannet nocturnal activity as a proportion of daylight hours | Proportion | The proportion of total nighttime hours that birds are expected to be active. |
| **Bird impacts - distributional responses** | | |
| Include distributional response for kittiwake | Yes/No | If this option is selected then displacement impacts will be predicted for kittiwake. Otherwise mortality predictions will be based on collision alone. |
| Mean peak seasonal densities | Birds per square kilometre | Estimated densities of birds in flight and on sea within the array area and displacement buffer (as defined in the total area of wind farm footprint parameter). |
| Displacement rates | Proportion | The proportion of birds expected to display distributional responses within the array area and displacement buffer. |

| | | |
|---|---|---|
| Seasonal displacement mortality rate | Proportion | The mortalities expected from distributional responses as a proportion of those birds displaying distributional responses. |
| **Bird populations** | | |
| Compare mortalities against a reference population | Yes/No | If this option is selected then mortalities are presented in the context of a focal population |
| Exclude impacts to birds from other populations | Yes/No | If this option is selected then mortalities are apportioned to represent only those relevant to a population of interest based on a user-defined proportion (see parameters below) |
| Exclude impacts to juveniles from the comparison | Yes/No | If this option is selected then mortalities are apportioned to represent only adult birds based on a user-defined proportion (see parameters below) |
| Exclude impacts to sabbatical birds from the comparison | Yes/No | If this option is selected then mortalities are apportioned to represent only birds breeding in any given year based on a user-defined proportion (see parameters below) |
| Reference population size | Individuals | The size of the population to which predicted mortalities should be compared. This is often defined as an adult breeding population of interest |
| Seasonal ratio of reference population to total population | Proportion | The proportion of mortalities expected to be attributable to the reference population. |
| Seasonal correction to be applied for adult ratio | Proportion | The proportion of mortalities expected to be adult birds. |
| Seasonal correction to be applied for sabbatical birds | Proportion | The proportion of mortalities expected to be non-sabbatical birds (i.e. the complement of the sabbatical rate). |

### 2 Default parameters

A number of parameters required for the analysis carried out by OCcAM have been included within the tool as default parameters. The choice of whether a required parameter should be user-defined versus defined within the tool was made based on judgements about those parameters expected to change from analysis to analysis, such as bird densities and turbine numbers and sizes, and also those parameters around which it was judged that the user may wish to deviate from the values that are currently widely applied for those parameter. Parameters selected to have default values within the tool were those that are difficult to obtain and therefore not expected to be readily available for indicative analyses, or those which are understood with a reasonable degree of certainty (for example, bird biometric parameters).

All default parameters used within the tool, and their sources, are displayed in the default parameters tab. This tab is locked to avoid accidental editing, but the sheet can be unlocked, allowing these values to be over-written. With the exception of bird flight height distributions, these data are directly used in calculations, so over-writing them will update outputs appropriately.

A list of the default parameters applied within OCcAM and their sources are listed in Table 2.

### 3 Calculation tabs

OCcAM includes several calculation tabs which are used to implement the collision risk modelling within the tool. These are adapted from those within the Band (2012) spreadsheet (BTO, 2013). The hours of daylight tab calculates daylight hours per day based on the user-input latitude and the algorithm defined in Forsythe *et al.* (1995). This is then summarised by month and used to determine monthly active hours for each species. The proportion at collision height tab uses the default bird flight height distributions data alongside the user-defined air gap and rotor diameter to calculate the proportion of birds at rotor height for each CRM species and parameter set. The single transit collision risk tab uses user input wind farm parameters alongside default bird biometric and flight speed parameters to determine the probability that a bird passing through the area swept by a rotating turbine blade will collide with that blade in the absence of any avoidance action. Finally, the overall collision risk tab brings together outputs from the previous three tabs as well as user-defined and default parameters to calculate the final collision mortality predictions according to the methodology described in Band (2012).

*Table 2 Default parameters provided within OCcAM, their sources and their application*

| Parameter set | Parameters included | Use within the tool | Sources |
|---|---|---|---|
| **Bird parameters** | | | |
| Biometric parameters | Average length, average wingspan, average flight speed, flight type and proportion of flights upwind for kittiwake and gannet. | Collision risk modelling - overall collision risk and single transit collision probability tabs | NatureScot (2023) |
| Bird flight height distributions | Proportion of birds expected to fly in 1 m height bands above sea level for kittiwake and gannet | Duplicated in the Collision risk modelling - proportion at collision height tab | Johnston *et al.* (2014) |
| Seasonal definitions | Season that each calendar month falls within for kittiwake and gannet | Collision risk modelling - overall collision risk tab | Furness (2015) |
| Apportioning values | Proportion of the population that is adult and sabbatical rate for all species | Apportioning of impacts - Results tab. May also be user-defined. | Proportion adult from stable age structure calculated from demographic parameters in Horswill and Robinson (2015); sabbatical rate taken from recent project EIAs e.g. Green Volt (2023) |
| **Wind farm parameters** | | | |
| Wind turbine specifications | Rotation speed, maximum blade width, average pitch and number of blades are provided for three turbine diameter ranges (<220 m, 220 - 250 m and >250 m) | Collision risk modelling - overall collision risk and single transit collision probability tabs | Average values derived from a proprietary database containing parameters used in avian collision risk modelling |
| Turbine operational time | Proportion of time turbines are expected to be operational based on wind availability and proportion of time turbines are likely to be stationary due to maintenance. | Collision risk modelling - overall collision risk tab | Default values provided in Caneco *et al.* (2022) |

### 4 Results tab and run log

Predicted impacts from collision and distributional responses, together with apportioning of mortality to the focal population (where required) are presented in the Results tab. Here, predicted mortalities are presented as total annual mortalities per parameter set and in total if required, and also broken down into each impact and biologically relevant season. In addition, if the option to compare to a reference population has been selected, mortality apportioned to the reference population will also be presented as a total, as a percentage of the refence population, and broken down by season and impact type.

An additional run log tab has been included in the tool to keep track of parameters used for a run and to document the rationale behind each selection.

### 5 Spreadsheet protection

Each sheet in the tool is locked such that only cells requiring user inputs can be edited. The data validation tools available within Excel have been used to ensure that invalid values cannot be entered into these cells. However, the sheets can be unlocked, allowing default parameters to be updated and other modifications to be made by the user if desired.

### 6 References


Band, B., 2012. Using a collision risk model to assess bird collision risks for offshore windfarms. Report by British Trust for Ornithology (BTO).

BTO, 2013. SOSS Collision Risk Modelling Excel Spreadsheet [Excel file].

Caneco, B., Humphries, G., Cook, A.S.C.P., Masden, E.A., 2022. Estimating bird collisions at offshore windfarms with stochLAB.

Forsythe, W.C., Rykiel, E.J., Stahl, R.S., Wu, H., Schoolfield, R.M., 1995. A model comparison for daylength as a function of latitude and day of year. Ecol. Model. 80, 87–95. https://doi.org/10.1016/0304-3800(94)00034-F

Furness, R.W., 2015. Non-breeding season populations of seabirds in UK waters; Population sizes for Biologically Defined Minimum Population Scales (BDMPS). (Natural England Commissioned Reports No. Number 164).

Green Volt, 2023. Green Volt Offshore Windfarm: Report to Inform Appropriate Assessment.

Horswill, C., Robinson, R.A., 2015. Review of seabird demographic rates and density dependence. (JNCC Report No. No: 552). Joint Nature Conservation Committee, Peterborough.

Johnston, A., Cook, A.S.C.P., Wright, L.J., Humphreys, E.M., Burton, N.H.K., 2014. Modelling flight heights of marine birds to more accurately assess collision risk with offshore wind turbines. J. Appl. Ecol. 31–41.

NatureScot, 2023. Guidance Note 7: Guidance to support Offshore Wind Applications: Marine Ornithology - Advice for assessing collision risk of marine birds.
